# Targeting AURKA by CRISPR/SaCas9 Alters Cell Cycle Progression and Enhances Lenvatinib-Associated Reduction in Cell Viability in Hepatocellular Carcinoma: An Integrated In Silico and HepG2 in Vitro Study

**DOI:** 10.64898/2026.11.17.767223

**Authors:** Noha Samir Taibe, Bangly Soliman, Elham Abdel-Badiea Mahmoud, Mahmoud El Hefnawi, Rania Hassan Mohamed

**Author notes:** corresponding author: Mahmoud Elhefnawi, Rania Hassan Mohamed. equal contribution (Mahmoud El Hefnawi, Rania Hassan Mohamed). Authors’ information: Noha Samir Taibe, Bangly Soliman, Elham Abdel-Badiea Mahmoud.

## Abstract

Hepatocellular carcinoma (HCC) presents substantial clinical concerns as a result of its high treatment resistance, frequent recurrence, and strong correlation with chronic liver diseases. Aurora kinase A (AURKA) is overexpressed in HCC correlating with poor prognosis and treatment resistance. Our bioinformatics analyses revealed that AURKA overexpression in HCC is associated with mitotic proteins and oncogenes, emphasizing its role in cancer progression. CRISPR/SaCas9-mediated AURKA knockdown markedly reduced AURKA expression levels through inducing several indels in the AURKA gene sequence, resulting in functional consequences including upregulation of p21, p53, and caspase-8, as well as downregulation of CDK1, CDK2, PI3K (alpha subunit), P38, and BCL-2 genes. Flow cytometry verified that these alterations were compatible with G2-phase cell cycle arrest and slight activation of apoptosis. Also, scratch and colony formation assays indicated a significant reduction in metastatic and proliferative potential. Furthermore, AURKA knockdown combined with Lenvatinib significantly reduce HepG2 cell viability compared with either treatment alone. Mechanistically, this study indicated that AURKA suppression reduced cell viability/proliferation by the modulation of oncogenic PI3K/AKT and p38/MAPK pathways. These *in-silico* and *in-vitro* findings highlight the critical role of AURKA in HCC pathogenesis. Here, the CRISPR/SaCas9-mediated genetic disruption of AURKA reveals its mechanistic role in regulating cell cycle, apoptosis, and tumor-suppressor signalling. Our study demonstrates AURKA as a potential molecular target to suppress HCC, while providing genetic evidence that its downregulation enhances Lenvatinib effect.

## Introduction

### Background

Worldwide, liver cancer ranks third in terms of cancer-related mortality and sixth in terms of cancer incidence [1]. Hepatocellular carcinoma (HCC) is a significant worldwide health issue, accounting for approximately 90% of primary liver malignancies [2]. In Egypt, where HCC is the most common malignancy in both sexes, the situation accompanying the HCC epidemic is very distinctive; more than 90% of all HCC cases in Egypt are caused by the high incidence of HCV[3, 4], which also greatly raises the nation’s HCC indices [5].

HCC still has a high overall morbidity and mortality rate, an aggressive growth habit, a high recurrence rate, and metastases are the primary cause of death for the majority of HCC patients[6]. Surgical resection, liver transplantation, local radiation, and systemic chemotherapy are among the treatment techniques that have been used to treat HCC in recent years [7]. Patients with HCC continue to have a dismal prognosis despite tremendous advancements in recent years in its treatment[8]. Patients with advanced liver cancer still have an inadequate prognosis, despite recent major advancements in targeted therapy, immunotherapy, interventional therapy, and radiation [9]. Numerous signalling pathways are known to be involved in the pathophysiology of HCC; however, it is yet unknown what molecular processes underlie the cancer’s malignant development [10]. Finding new targets for customized treatments and more potent biomarkers for early detection is, therefore, extremely important from a therapeutic standpoint [11]. For this reason, finding new genes for HCC diagnosis and precision medication is crucial [12].

Aurora kinase A (AURKA) is a mitotic serine/threonine kinase family member that is vital for controlling the cell cycle and mitosis (centrosome maturation and separation, spindle assembly, chromosome alignment, and G2 to M transition), both of which are necessary for preserving the integrity of genetic material [13, 14]. According to The Cancer Genome Atlas (TCGA) UALCAN database, Aurora kinases are expressed in a wide range of cancers such as hepatocellular carcinoma (HCC) [15], breast cancer[16–18], gastric cancer [14], lung cancer [19, 20], and thyroid carcinoma [21]. AURKA functions as an oncogene by regulating multiple biological processes involved in the regulation of cancer cell proliferation, migration, invasion, epithelial-mesenchymal transition (EMT), apoptosis, metastasis, genomic instability, and cancer stem cell (CSC) behaviours. All of which lead to acceleration of cell cycle progression and promotion of tumor metastasis by activating several signalling pathways and consequently the development of tumours [15]. Further research on the function of AURKA in HCC and its underlying mechanisms is necessary in light of the possibility that AURKA could be used as a therapeutic target for the management of HCC[22–24].

The RNA-guided nuclease linked with CRISPR (clustered regularly interspaced short palindrome repeats) Cas9 can be efficiently introduced at particular locations in the genome for targeted loss-of-function mutations [25]. The two primary parts of this system are single-guide RNA (sgRNA) and Cas9 endonuclease. Streptococcus pyogenes Cas9 (SpCas9) or Staphylococcus aureus Cas9 (SaCas9) are currently the most frequently employed Cas9 proteins, SaCas9(about 3.1 kb) is smaller than SpCas9 (about 4.7 kb) and comprises just 1053 amino acids, making it simpler to deliver to the target cells[26]. SaCas9 could boost editing effectiveness when the length of the guide sequence increased from 20 to 21⁓23 nt. Therefore, SaCas9 is better suited for therapeutic use [27].

In this study, AURKA was selected using bioinformatics integrative analysis that has been previously published [28–31]. Our purpose in this study is to use the CRISPR/SaCas9 system to knockdown the AURKA gene in HepG2 cells and evaluate HepG2 cells’ subsequent tumor regression using our experimental procedure. As illustrated in Figure 1, the effectiveness of CRISPR/SaCas9-mediated AURKA knockdown is assessed, and its effects on cell viability, proliferation, and migration are examined, along with its effects on the expression of important cell cycle and apoptosis regulators and related oncogenic signalling pathways.

**Figure 1:**
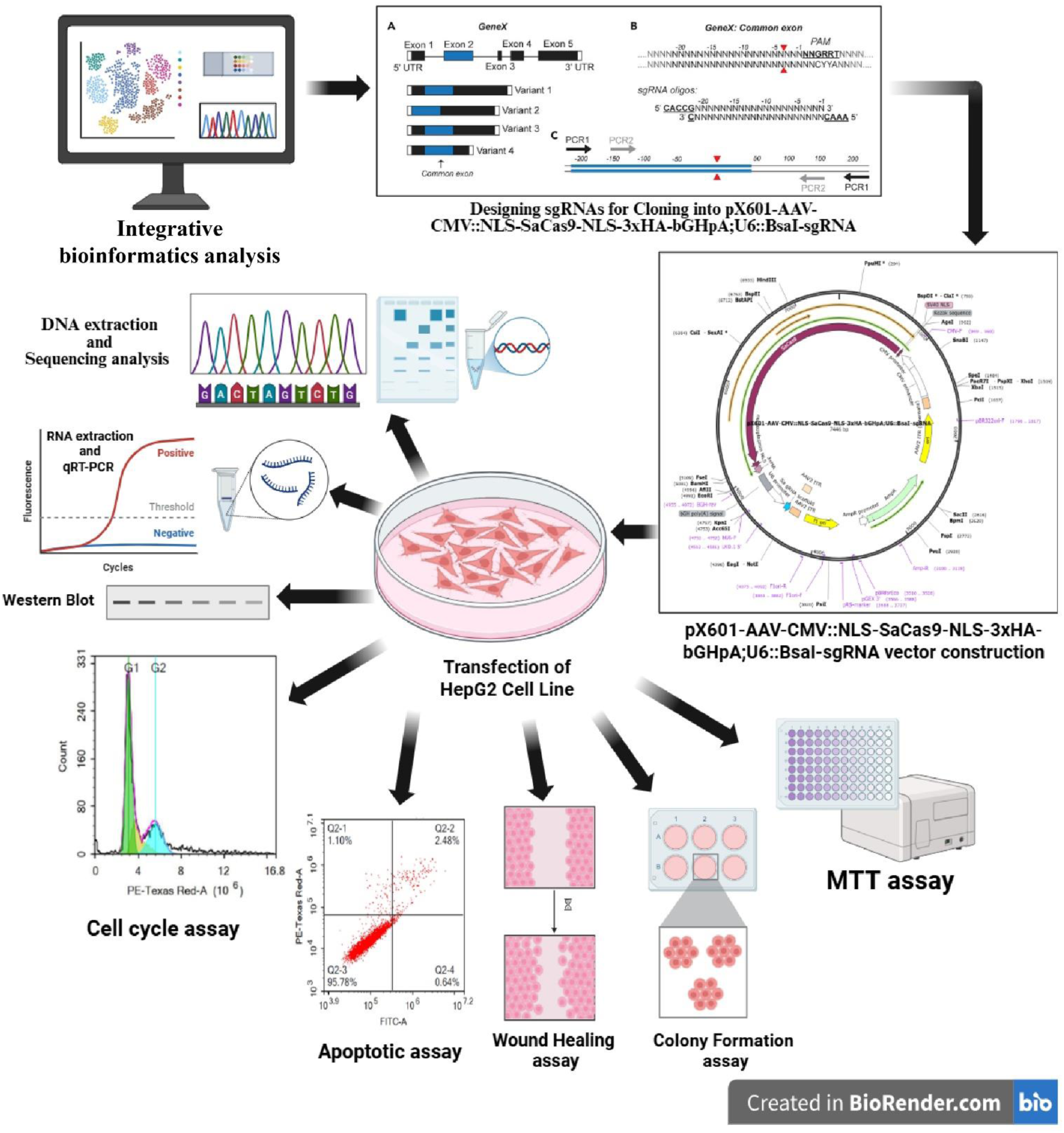
Schematic workflow of SaCas9-mediated AURKA gene editing in human hepatocellular carcinoma (HCC) cells. Integrative bioinformatics analysis was done on human HCC datasets to predict the expression and role of AURKA. AURKA-specific single guide RNAs (sgRNAs) were designed and cloned into the pX601-AAV-CMV::NLS-SaCas9-NLS-3xHA-bGHpA;U6::BsaI-sgRNA vector. Then, HepG2 cells were transfected with the recombinant and null vectors. Several downstream investigations, such as sequencing analysis, RT-PCR, and Western blotting were carried out after transfection to assess the transfection and the impact of gene editing. Functional assays such as MTT, colony formation, wound healing, apoptotic, and cell cycle assays were performed to evaluate the effect of AURKA downregulation on the treated HepG2 cells.

## Methods

### Integrative Bioinformatics Analysis

The AURKA gene was selected based on findings from previous studies [28, 29]. To evaluate its differential expression, multiple R packages were utilized following a previously published protocol [32], along with the Gene Expression Profiling Interactive Analysis (GEPIA) platform [33]. AURKA expression was examined at both the mRNA and protein levels.

For the RNA-seq analysis, RNA-seq expression data (RSEM expected counts) were obtained from the UCSC Xena Toil Recompute Compendium (“TCGA_Target_GTEx_gene_expected_count” dataset). Differentially expressed genes (DEGs) were identified between GTEx normal liver and TCGA Liver Hepatocellular Carcinoma (LIHC) samples using thresholds of |log2FC| ≥ 1.5 and Benjamini–Hochberg adjusted p-value < 0.05 [34]. To identify outliers, the top 75% of the genes were filtered and then clustered.To identify modules (clusters) of highly co-expressed genes, Weighted Gene Co-expression Network Analysis (WGCNA) was performed using the WGCNA R package. Several soft-thresholding powers (β) were assessed to guarantee a scale-free topology, which is a basic WGCNA principle. A power of β = 10 was chosen using the scale-free topology fit index.

Genes from the AURKA-associated module were further analyzed for prognostic significance using the survival and survminer packages for pathway enrichment through over-representation analysis (ORA) with clusterProfiler, referencing hallmark gene sets from MSigDB. The intersection of DEGs, hallmark gene sets (the G2M checkpoint), and top survival genes was visualized in a heatmap [35, 36].

To study the link between AURKA and immune invading cells such as T Cell Cytotoxic (CD8A, PRF1, GZMB), B Cell (CD79A, CD19, MS4A1) Macrophage (CD68, CD163, ITGAM), Dendritic Cell (FLT3L, CLEC9A, LAMP3), the R package “GSVA” [37] was used in conjunction with single-sample Gene Set Enrichment Analysis and the R package “ESTIMATE” [38] was used to analyse the correlation between AURKA expression and stromal non-tumor /immune cell infiltration across the tumor microenvironment (TME) of LIHC. Spearman correlation analysis was conducted to assess the relationship between AURKA expression, immune invading cells, and immune checkpoint molecules (inhibitors such as PDCD1, CTLA4, TIGIT, LAG3, HAVCR2, CD274, and co-stimulators such as ICOS, CD28, TNFRSF9, TNFRSF18). Spearman’s correlation coefficient (ρ) was considered significant at p < 0.05. Also, the functional impact of *AURKA* expression on predicted immunotherapy outcomes was assessed using the Tumor Immune Dysfunction and Exclusion (TIDE) (http://tide.dfci.harvard.edu) tool. All LIHC samples were assigned a TIDE score, which predicts patient response to Immune Checkpoint Blockade (ICB), with a larger score indicating a higher risk of resistance.

Beyond transcriptional analysis, AURKA protein expression in LIHC was also evaluated using UALCAN (http://ualcan.path.uab.edu) and The Human Protein Atlas (www.proteinatlas.org).

### sgRNA design and plasmid construction

pX601-AAV-CMV::NLS-SaCas9-NLS-3xHA-bGHpA;U6::BsaI-sgRNA vector (Addgene plasmid # 61591)[27] was purchased from Addgene and preserved at −80 °C. AURKA gene (Gene ID: 6790, NCBI) has 13 transcripts; we selected the common exon among them to generate sgRNAs using CRISPOR [39]. Two sgRNAs were selected as they showed the highest specificity and efficiency scores that have no perfect matches (0 mismatches) elsewhere in the genome and show minimal potential for off-target binding (≤2 mismatches in non-critical regions) as predicted by CRISPOR [39] and CHOPCHOP tools [40] [41].

According to [42], as shown in Table 1, forward and reverse oligos for each sgRNA were designed and then synthesized by Macrogen, USA. The two sgRNAs were inserted into the vector according to [43] with some modifications. The vector was digested with Bpil (BbsI) enzyme (Thermo Scientific) and subsequently dephosphorylated by Invitrogen’s calf intestinal alkaline phosphatase (CIAP). ThermoScientific’s GenJet gel extraction kit was used to purify the linear dephosphorylated vector following dephosphorylation.

**Table 1:**
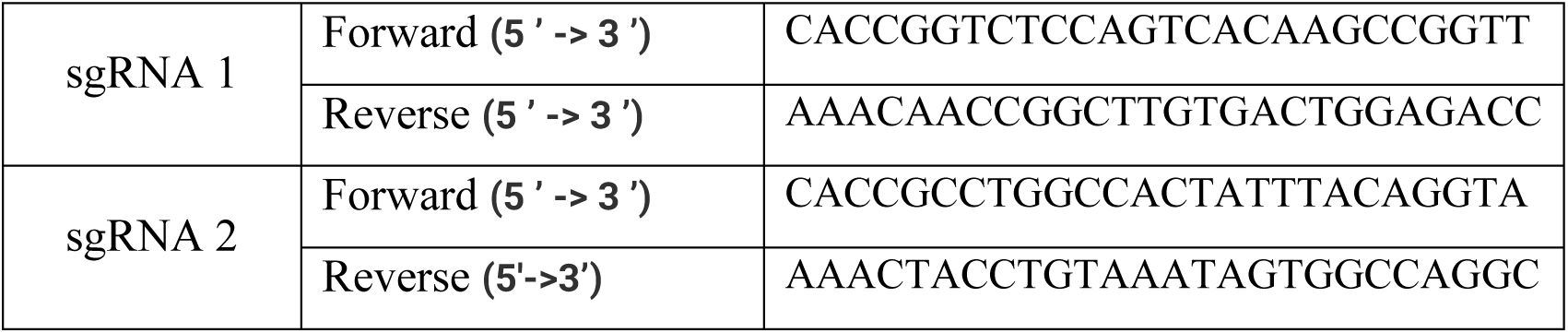
Sequences of sgRNA.

Then each pair of oligos was annealed and phosphorylated using T4 PNK(ThermoScientific)[43]. The reactions were conducted by a SimpliAmp thermal cycler (Applied Biosystems) with the following conditions:37°C for 30 min, followed by 95°C for 5 min, and then ramped down to 25°C at 5°C/min. The two phosphorylated and annealed oligos were diluted (1: 200), then ligated to the digested dephosphorylated vector (pX601-AAV-CMV::NLS-SaCas9-NLS-3xHA-bGHpA;U6::BsaI-sgRNA) withT4 DNA ligase (Thermo Scientific), according to the manufacturer’s instructions. The recombinant plasmids were transformed into competent HB101 cells according to the Addgene Bacterial Transformation protocol. Colony PCR was used for screening and selecting the transformed colonies. Plasmid DNA purification was performed according to the protocol of the PureLink™ Expi Endotoxin-Free Maxi Plasmid Purification Kit (Invitrogen) for sequencing using the U6 Universal Primer, as shown in Table 2, by Humanizing Genomics Macrogen (Korea).

**Table 2:**
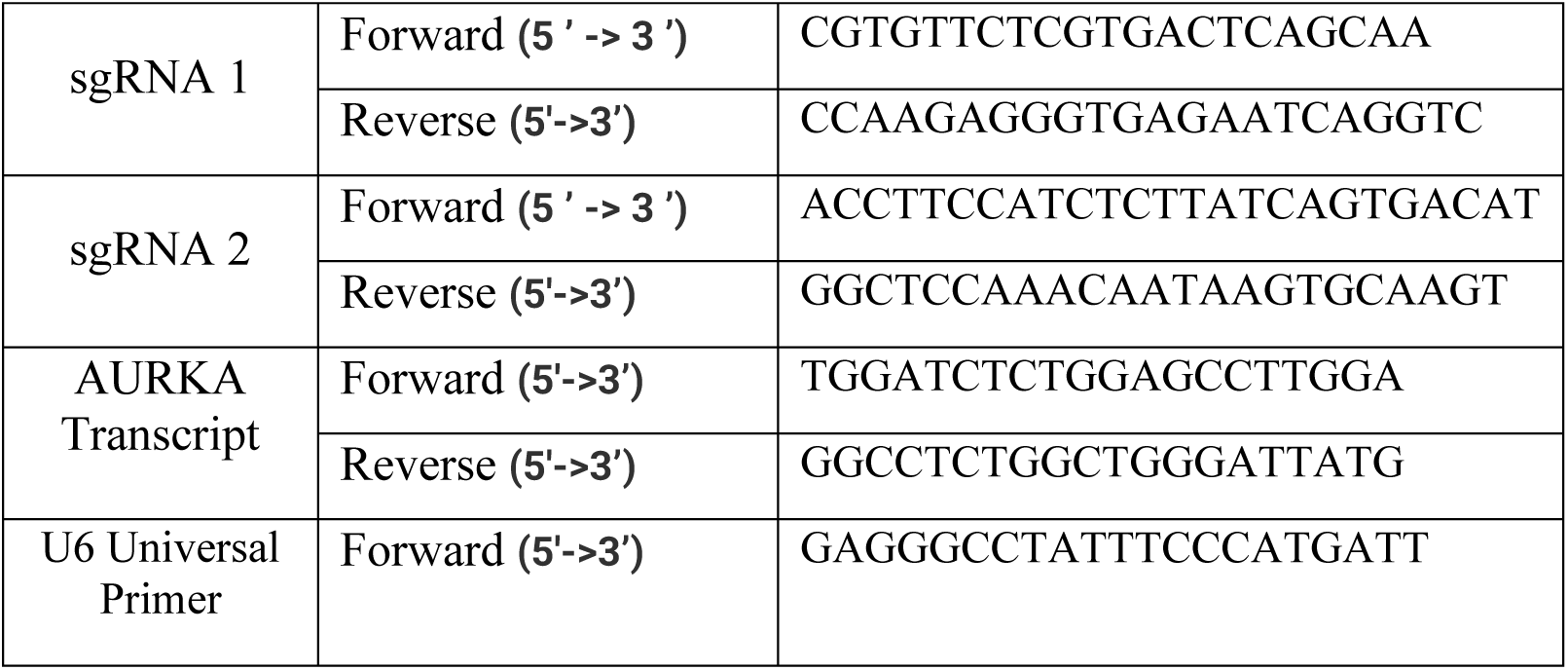
Primers for genomic analysis, sequencing, and qRT-PCR.

### Cell Culture and Transfection

HepG2 (Human hepatocellular cancer) cells were obtained from the International Centre for Training and Advanced Researches (ICTAR-Egypt). The cells were cultured in RPMI 1640 medium (Gibco, USA) containing 10% heat-inactivated FBS (fetal bovine serum, Gibco), 1% penicillin/streptomycin (Sigma, USA) at 37 °C, and 5% CO_2_ until 70-90 % confluence. About 1.5×10^5^ (cells/well) of HepG2 cells were seeded in 6-well plates. Cell cultures were transfected according to the Lipofectamine 3000 transfection agent protocol (Invitrogen, USA) with a null vector, gRNA1, gRNA2, and both gRNA1 and gRNA2(gRNA1+gRNA2). The transfection efficiency was confirmed by Western blot using an anti-HA antibody in comparison to untreated cells. Following the transfection, the editing efficiency was confirmed by sequencing, RT-PCR, and Western Blot assay.

### Sanger Sequencing

After transfection, genomic DNA was extracted from transfected cells after 72 hours. PCR amplification was performed using the AmpliTaq Gold® 360 Master Mix (Invitrogen) protocol; the primers used are presented in Table 2. The PCR products were loaded on a 1% agarose gel and then purified using the GeneJET Gel Extraction Kit (Thermo Scientific). Purified PCR amplicons were sequenced with the previous primers by Humanizing Genomics Macrogen (Korea). MUSCLE alignment was used to analyse the results of sequencing, and the alignment outputs were visualized using Jalview software [44]

### RNA Extraction and Quantitative RT-PCR

Total RNA was extracted from transfected cells after 72 hours using the TRIzol™ Reagent (Invitrogen, USA) protocol. The purity and concentration of RNA were confirmed using the Qubit RNA IQ Assay Kit by Qubit 4 Fluorometer (Invitrogen, USA). cDNA was synthesized from total RNA using the RevertAid First Strand cDNA Synthesis kit protocol. The quality and quantity of cDNAs were measured using the Qubit dsDNA HS Assay Kit through Qubit 4 Fluorometer. The real-time RT-PCR reactions were done using primers targeting the following genes: AURKA (as shown in Table 2), CDK1, Caspase 8, BCL-2, P53[45], P21[46], CDK2, CDC, P38, PI3K(alpha subunit), and the human GAPDH (Glyceraldehyde 3-phosphate dehydrogenase) as a control (reference gene) according to Maxima SYBER Green qPCR Master Mix(2x), no ROX (Thermo Scientific) protocol by a thermal cycler (Applied Biosystem 7500 Fast) with the following conditions: 1 cycle of initial denaturation at 95°C for 10 min, 40 cycles of PCR amplification (Denaturing at 95°C for 15 s, Annealing/Extending at 60°C for 60 s). Relative mRNA expression was calculated using the 2^−ΔΔCt^ method[47].

### Western Blot

Total proteins were extracted from transfected cells after 72 hoursusing ice-cold RIPA lysis buffer (150 mM NaCl, 0.5 mM Tris HCl pH 8.0, Triton-X100, 1.0% Sodium deoxycholate (Thermo Scientific), 0.1% SDS,10% glycerol) and freshly added proteases inhibitors (1μl from PMSF (100mM), 5μl from EDTA (100mM), 1μl from NaF (100mM), 1μl from Na_3_VO_4_ (100mM, Sigma)). The proteins were quantified using the Qubit Protein BR Assay Kit, and 25μg of protein was loaded in a 15% SDS-PAGE gel, and then transferred to the PVDF membrane (Thermo Scientific) using the XCell II™ Blot Module protocol (Invitrogen, USA). Following the transfer of proteins, MemCode™ Reversible Protein Stain Kit was used for staining the membranes. After destaining the membranes, the membranes were incubated overnight at 4°C with primary antibodies such as anti-Aurora A Rabbit monoclonal antibody 1:5000 [EPR5026, abcam], CDK1 Recombinant Rabbit Monoclonal Antibody 1:1500 (SM01-44, Invitrogen), anti-HA mouse monoclonal antibody 1:4000 (HT301, TRANS), and GAPDH mouse monoclonal antibody 1:4000 (3E8AD9, Invitrogen). Then, the membranes were incubated with secondary antibodies Goat anti-Rabbit IgG (H+L) Secondary Antibody, HRP 1:10,000 (31460, Invitrogen) or Goat anti-Mouse IgG (H+L) Secondary Antibody, HRP 1: 100 (32430, Invitrogen) for 2 hours at 16°C. The proteins were detected according to the SuperSignal West Atto Ultimate Sensitivity Chemiluminescent Substrate protocol (A38555, Thermo Scientific). The images of the membranes were captured with the ChemiDoc XRS imaging system (Bio-Rad, USA). The intensity of protein bands was measured using ImageJ software[48].

### Cell Cycle Assay

At 72 h post-transfection, the cells were collected by trypsinization and washed twice with ice-cold PBS. Cells were resuspended in 2 mL of 60% ice-cold ethanol and incubated at 4°C for 1 h for fixation. Fixed cells were washed twice with PBS and resuspended in 1 mL of PBS containing 50 μg/mL RNAase A and 10 μg/mL propidium iodide (PI). After 20 min of incubation in the dark at 37 C, cells were analyzed for DNA content by flow cytometry analysis (12,000 events were acquired) using the FL2 (λex/em 535/617 nm) signal detector (ACEA Novocyte™ flow cytometer, ACEA Biosciences Inc., San Diego, CA, USA) [49–54].

### Apoptotic Assay

Apoptotic and necrotic cell populations were determined using the Annexin V-FITC apoptosis detection kit (Abcam Inc., Cambridge Science Park, Cambridge, UK) coupled with 2 fluorescent channels of flow cytometry (12,000 events were acquired). At 72 h post-transfection, cells were collected by trypsinization and washed twice with ice-cold PBS. Then, cells were incubated with 0.5 ml of Annexin V-FITC/PI solution for 30 min in the dark at room temperature, according to the manufacturer’s protocol. After staining, cells were injected via ACEA Novocyte™ flow cytometer (ACEA Biosciences Inc., San Diego, CA, USA) and analyzed for FITC and PI fluorescent signals using FL1 and FL2 signal detectors, respectively (λex/em 488/530 nm for FITC and λex/em 535/617 nm for PI). Positive FITC and/or PI cells were quantified by quadrant analysis and calculated using ACEA NovoExpress™ software (ACEA Biosciences Inc., San Diego, CA, USA)[49, 50, 53, 54].

### Wound Healing Assay

At 72 h post-transfection and at a cell confluency of 90%, a uniform scratch wound was made on the cell monolayer using a 100 µl pipette tip. After removing the media and washing the plates three times with PBS, fresh media was added to the cell culture plates, and photographs of the scratch width at three distinct positions: bottom, middle, and lower were taken immediately (zero time) by an inverted microscope (Carl Zeiss 204 Microscopy GmBh, Germany). At 24, 48, and 72 hours, the width of the scratch was tracked and recorded. ImageJ software[48] was used to quantify the wound areas, and the assay was performed in triplicate independent assays.

### Colony Formation Assay

Transfected HepG2 cells (3×10^3^ cells/well) were cultured in 6-well plates for 14 days at 37°C and 5% CO_2_, and the cells transfected with the null vector were used as a control. After the incubation, the media were removed, and the colonies were rinsed with PBS twice and fixed with 2 mL of 1 acetic acid:7 methanol at room temperature for 5 min. After washing three times with PBS, the colonies were stained with 400 μl of 0.5% crystal violet for 30 min at room temperature. After incubation, the plate was washed with tap water and left to dry at room temperature[55]. The colonies were counted and imaged using an inverted microscope and quantified using ImageJ [48].

### Cell cytotoxicity

HepG2 cells (8×10^3^) were seeded in a 96-well plate at 37°C and 5% CO_2_ for 24 hours until 70% confluency; then the media was aspirated carefully, and the cells were transfected with both gRNA1 and gRNA2 (gRNA1+gRNA2) using Lipofectamine 3000 protocol and, where indicated, 5 μM Lenvatinib (Sigma Aldrich) was added. Null-transfected HepG2 cells were used as a negative control, while HepG2 cells treated with 5μM Lenvatinib drug only were used as a positive control.

The transfected cells were incubated for 48 hr, and then the media were removed without disturbance, and 25 μl of 5 mg/ml MTT solution (Sigma Aldrich) was added per well, and the cells were incubated in a dark place for 3 hours until crystal formation at 37°C and 5% CO_2_. 100 μL of DMSO (dimethyl sulfoxide) was added per well, and then the plate was incubated for 15 minutes at room temperature under microscopic monitoring until the crystals dissolving. Cell viability was evaluated using a microplate spectrophotometer (sunrise TECAN) to measure absorbance values at 570 nm. The cell viability percentage was calculated through this equation: Cell viability % = (average of O.D_570_ of sample/average of O.D_570_ of negative control) *100 [56].

The enhanced effect of the combination of Lenvatinib with both gRNA1 and gRNA2 (gRNA1+gRNA2) was calculated using the Bliss score equation [57] as follows:

Step 1: Fractional inhibition “F”

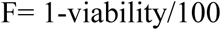

Step 2: Bliss expected effect “Bliss exp”

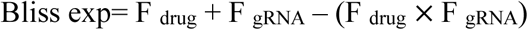

Step 3: Bliss score “S”

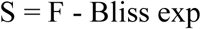

Bliss scores greater than zero indicate enhanced interaction, less than zero indicates antagonistic interaction, while equal to zero indicates additive interaction [58].

### Statistical analysis

All experiments were performed as three separate experiments. GraphPad Prism version 8.0.2 was used to code, tabulate, and statistically analyse the collected data using unpaired t-tests or ordinary one-way ANOVA. Two-way ANOVA and the Tukey-Kramer multiple comparisons test were used in the statistical study to compare the differences between the groups. The data are presented as the mean ± standard deviation (SD). Significant differences are indicated by the symbols *p≤0.05, **p≤0.01, ***p≤0.001, ****p≤0.0001, and ns (not significant) p>0.05.

## Results

### AURKA is overexpressed in liver hepatocellular carcinoma (LIHC)

To investigate the expression of AURKA gene in hepatocellular carcinoma, we performed network analysis of the normal GTEx liver gene dataset and the TCGA LIHC dataset. After performing Principal Component Analysis (PCA), five samples were identified as outliers and eliminated prior to further analysis, as shown in supplementary Figure S1.

According to the volcano plot in Figure 2A, differential expression analysis on 358 TCGA-LIHC primary tumours and 107 GTEx normal liver tissue samples with |logFC| > 1.5 and adjusted p < 0.05 showed that the AURKA gene was among 661 genes that were significantly upregulated (red dots), 12967 genes that were not significant (grey dots), and 1616 genes that were significantly downregulated (blue dots). AURKA expression was upregulated in the TCGA LIHC dataset versus the normal GTEx liver gene dataset with a P value =1.75e-35, as illustrated in Figure 2B. Additionally, as shown in Figure 2C, the AURKA protein expression was significantly overexpressed in HCC samples (n=165) relative to normal samples (n=165). When assessing its correlation with HCC development, we found that AURKA gene expression is linked to a high mortality rate, as demonstrated by survival analysis in Figure 2D, which shows that patients with high expression of the gene passed away sooner than those with low expression of AURKA. Weighted Gene Co-expression Network Analysis (WGCNA) analysis, as shown in Figure 2E and supplementary Figure 2, revealed that 253 AURKA-Green module genes with log2FC ≥1.5 and p.adj <0.05 are significantly correlated with clinical traits of HCC, including pathological stage, histological status, and fibrosis.

**Figure 2:**
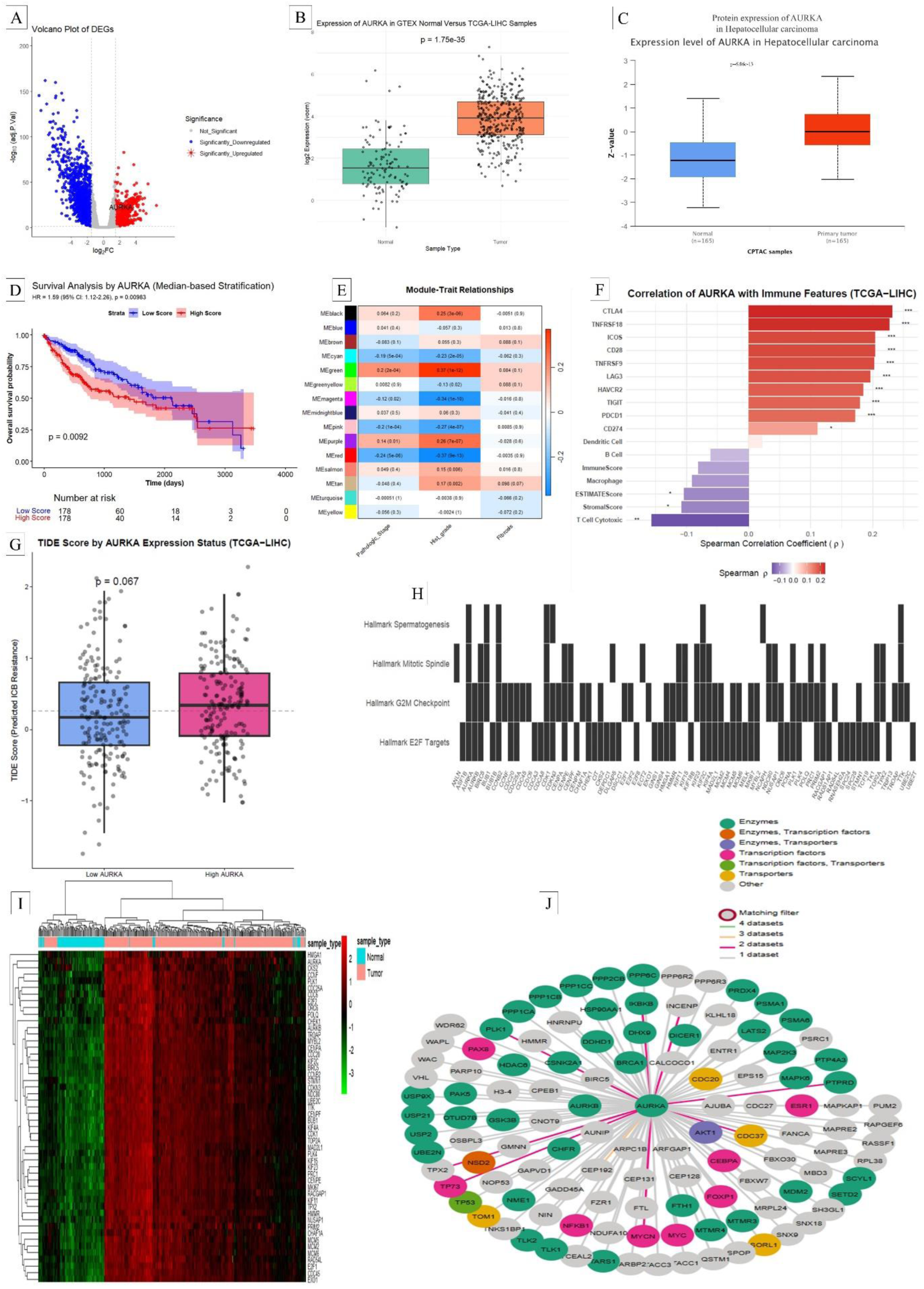
HCC cells exhibit overexpression of AURKA gene. A) Volcano plot of differential gene expression (DEG) analysis. B) Box plot represents the gene expression of AURKA in GTEX normal samples versus the TCGA-LIHC Samples. C) Protein expression of AURKA in Hepatocellular Carcinoma. D) Effect of AURKA expression level on the survival time of HCC patients. E) Gene set enrichment analysis of module traits related to pathological stage, histological grade, and fibrosis. F) AURKA correlation with Immune Features in TCGA-LIHC. G) In TCGA-LIHC, AURKA overexpression predicts resistance to immune checkpoint blockade (ICB). H) AURKA-related ORA (over-representation analysis) using MSigDB Hallmark gene sets as spermatogenesis, Mitotic spindle, E2F targets, and G2M checkpoint. I) Heatmap of G2M checkpoint genes. J) Biological General Repository for Interaction Datasets (BioGRID) chart of AURKA protein.

The comprehensive immune infiltration and predictive correlation analysis using ssGSEA and ESTIMATE scores revealed that AURKA expression shows positive correlation with multiple immune checkpoint molecules in TCGA-LIHC, including CTLA4, TNFRSF18, ICOS, CD28, and PDCD1 (PD-1), and a significant negative correlation with the infiltration levels of key anti-tumor immune cells, including cytotoxic T cells and with the ESTIMATE and stromal scores, as shown in Figure 2 F.

Tumors with high and low AURKA expression did not differ substantially, according to TIDE analysis (Wilcoxon rank-sum test, p = 0.067), as shown in Figure 2G and the TIDE score displayed in the supplemental table.

Using hallmark gene sets such as spermatogenesis, mitotic spindle, E2F targets, and G2M checkpoint, over-representation analysis (ORA) showed the expression pattern of the AURKA gene and its functionally associated genes in both normal and liver hepatocellular carcinoma (LIHC) samples, as shown in Figure 2 H. Furthermore, the heatmap (Figure 2G) displays the expression pattern of AURKA and its functionally associated genes in both normal cells and liver hepatocellular carcinoma (LIHC) at various stages. These findings suggest that, particularly in the early stages of HCC, as shown in supplementary Figure 3 E, AURKA is markedly overexpressed in HCC tumors in comparison to normal tissue. Normal tissue exhibits low levels of AURKA expression, while tumor stages, particularly Stage I and II, show markedly higher levels, suggesting early amplification during carcinogenesis. In parallel with AURKA, all cancer stages have increased expression of the CDK1, CDC20, CDK2, TPX2, PLK1, CHEK1, MYC, AKT1, MKI67, and BCL2 genes.

Furthermore, as Figure 2 J shows, AURKA plays an important role in cancer pathways by interacting with proteins and transcription factors involved in cell cycle regulation, DNA repair, and signal transduction. AURKA’s interactions with immune-related proteins such as STAT3 and TP53 indicate that it modulates immune responses in tumours. These findings emphasise AURKA’s involvement as a major hub in oncogenic signalling in hepatocellular carcinoma, influencing proliferation, genomic stability, and immune function.

These findings suggested that the higher expression of the AURKA gene was associated with tumor recurrence or resistance to treatment. Its role as a master regulator of mitotic progression and tumor cell proliferation is reinforced by its co-expression with important mitotic regulators (CDK1, PLK1, CDC20), oncogenes (MYC, AKT1), and proliferation markers (MKI67). Particularly in the early stages of HCC, AURKA is markedly overexpressed in HCC tumors in comparison to normal tissue.

### AURKA expression was downregulated with sgRNAs/SaCas9 genome editing

To investigate the efficiency of sgRNAs/SaCas9 genome editing to knockdown AURKA gene expression, we performed sequence analysis on the edited region, AURKA expression using RT-PCR, and Western blot analysis. Sequence analysis revealed that gRNA1 and gRNA2 showed several indels compared with the wild-type gene sequence, as shown in Figure 3A. These indels led to a significant decrease in the relative gene expression of AURKA gene. Using gRNA1 reduced AURKA expression to 0.34-fold with a p-value ≤0.01 in comparison to the control. Also, gRNA2 reduced AURKA expression to 0.2-fold in comparison to the control with a p-value ≤0.001. Moreover, the combination of both gRNAs (gRNA1+gRNA2) exhibited the lowest relative AURKA gene expression 0.1-fold in comparison to the control and both gRNAs with a p-value ≤0.001, as shown in Figure 3B.

**Figure 3:**
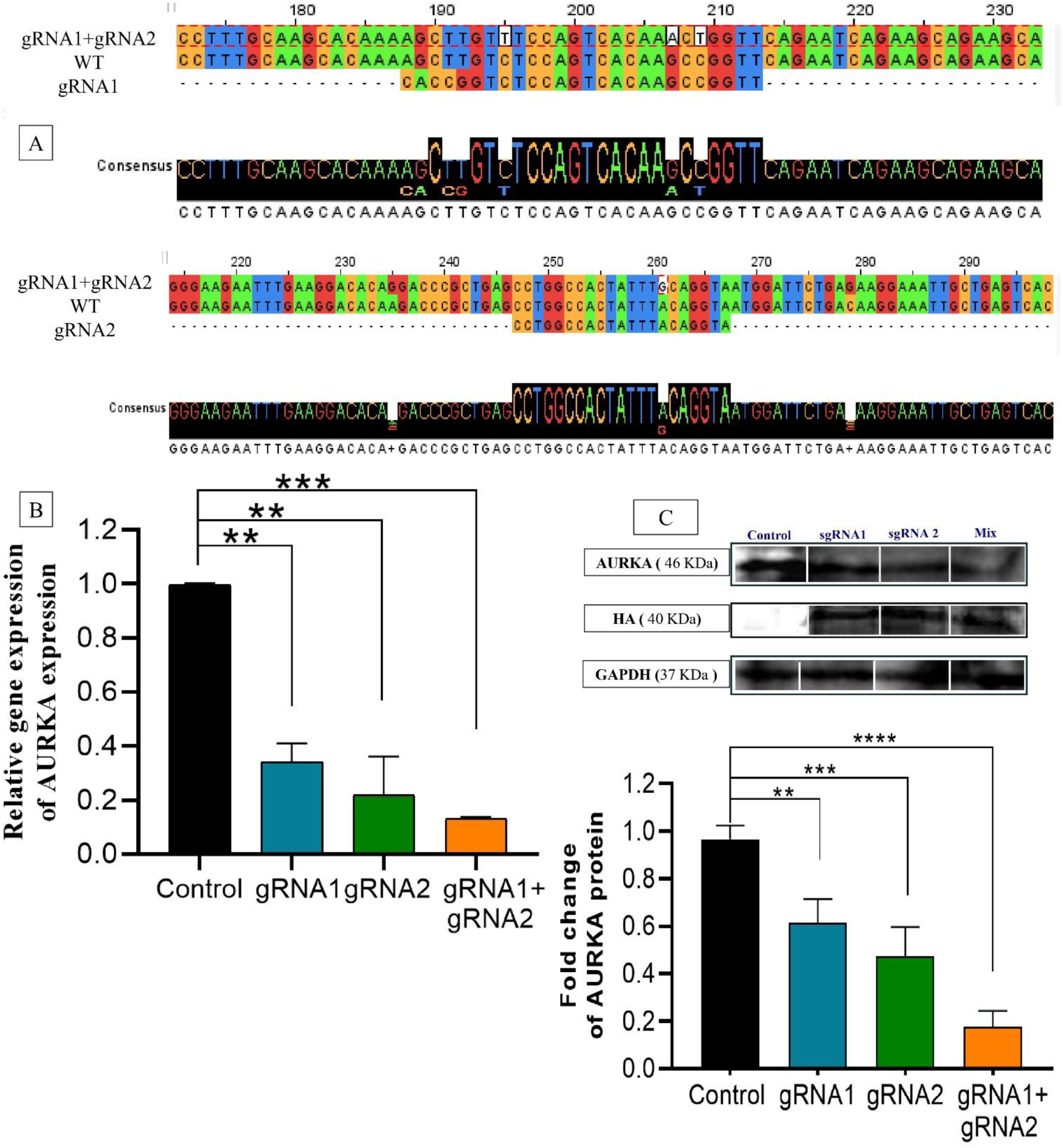
Downregulation of AURKA expression in HepG2 cells after gene editing. A) Analysis of the genome sequences of HepG2 cells treated with gRNA1+gRNA2 using the MUSCLE alignment tool, with Jalview visualisation. B) AURKA’s relative gene expression in HepG2 cells treated with a null vector (Control), HepG2 cells treated with gRNA1only (gRNA1), HepG2 cells treated with gRNA2 only (gRNA2), and HepG2 cells treated with both gRNA1 and gRNA2 (gRNA1+gRNA2). C) Western blot analysis of AURKA protein in Control, gRNA1, gRNA2, and gRNA1+gRNA2. All results were normalized to GAPDH. All experiments were performed as three independent experiments. GraphPad Prism version 8.0.2 was used to code, tabulate, and statistically analyse the collected data using ordinary one-way ANOVA. The data are presented as the mean ± standard deviation (SD). The significant difference is indicated by the symbols *p≤0.05, **p≤0.01, ***p≤0.001, ****p≤0.0001, and ns (not significant) p>0.05.

Figure 3C illustrates how the AURKA protein expression decreased in proportion to the decrease in gene expression. In comparison, gRNA2 reduced protein fold change by 0.47, and gRNA1 reduced protein fold change by 0.615, with p-values of ≤0.001 and ≤0.01, respectively. Using both gRNA1 and gRNA2 (gRNA1+gRNA2) demonstrated a significant reduction in AURKA protein expression with a 0.177 protein fold change (p-value ≤0.0001). As the combination of gRNA1 and gRNA2 treatment (gRNA1+gRNA2) had the strongest effect on AURKA gene knockdown, we decided to use this treatment mainly in the subsequent experiments. These results indicate the successful AURKA knockdown in HepG2 cells by CRISPR/SaCas9 treatment.

### AURKA knockdown induces cell cycle arrest in HepG2 cells

We investigated the effect of AURKA knockdown on the HepG2 cell behaviour and characteristics. Considering the role of AURKA in cell cycle regulation [59], we first assessed AURKA downregulation’s impact on the cell cycle and the key genes regulating it in HepG2 cells. Confirming the genes we found here associated with AURKA through our bioinformatics analysis (as shown in Figure 2).

In cells treated with gRNA1 and gRNA2, the relative gene expression of P21 was upregulated by 7.09-fold compared to the controlwith a p-value ≤0.01, as shown in Figure 4A. Moreover, CDK2, CDK1, and PI3K (alpha subunit) genes exhibited significant downregulation in relative gene expression of 0.13, 0.21, and 0.27-fold, respectively, as shown in Figure 4B-D, with a P-value ≤0.01 compared to the control.

**Figure 4:**
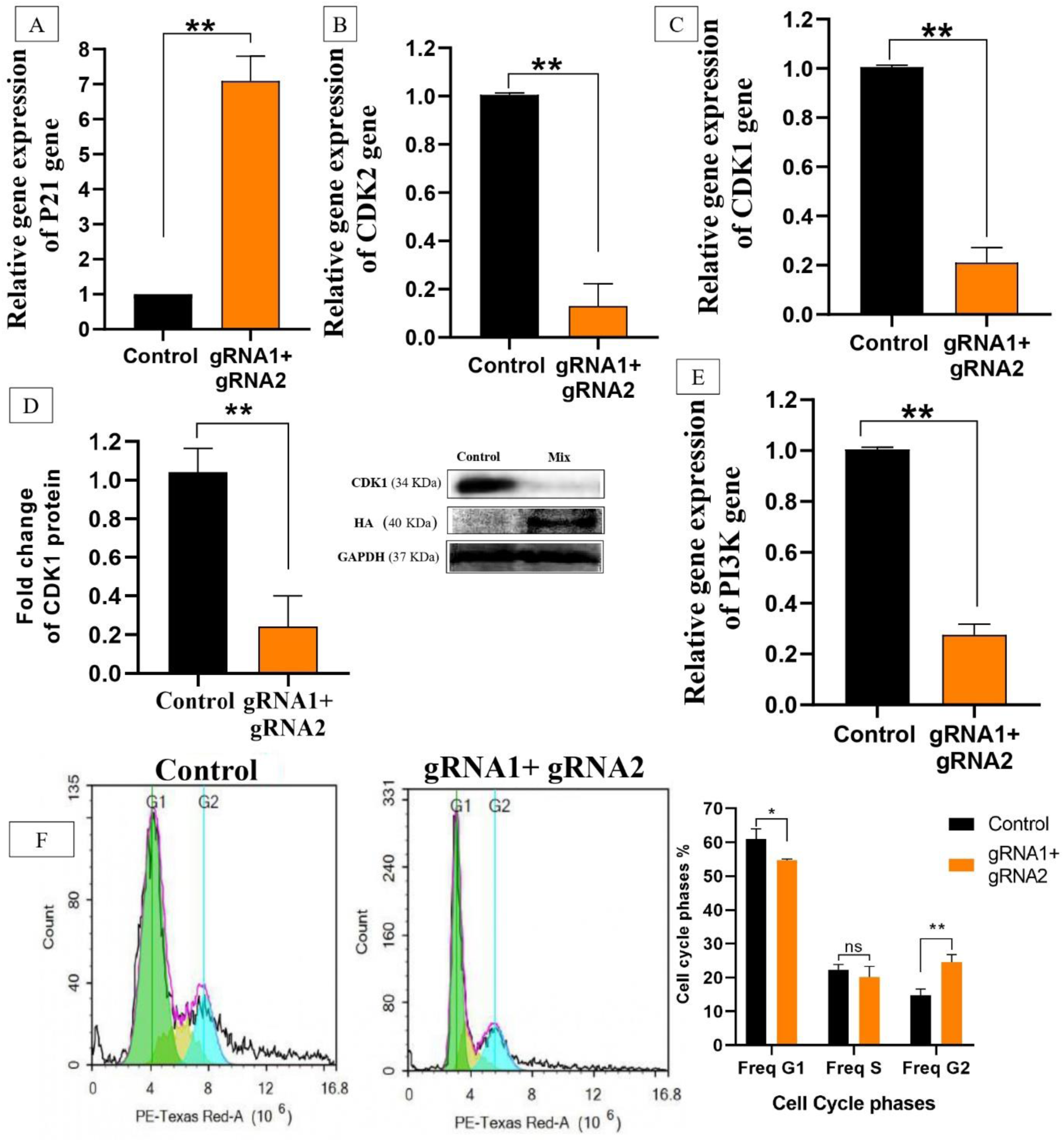
HepG2 cell cycle progression is suppressed by AURKA downregulation. AURKA downregulation impact on the relative expression of several cell cycle genes in Control (null vector) and gRNA1+gRNA2 (HepG2 cells treated with both gRNA1 and gRNA2): A) P21, B) CDK2, C) CDK1, E) PI3K (alpha subunit). D) Western blot analysis of the CDK1 gene in Control and gRNA1+gRNA2. F) Cell cycle analysis using flow cytometry in Control and gRNA1+gRNA2. All results were normalized to GAPDH. All experiments were performed as three independent experiments. GraphPad Prism version 8.0.2 was used to code, tabulate, and statistically analyse the collected data using unpaired t-tests or two-way ANOVA, and the Tukey-Kramer multiple comparisons test was used in the statistical study to compare the differences between the groups. The data are presented as the mean ± standard deviation (SD). Significant differences are indicated by the symbols *p≤0.05, **p≤0.01, ***p≤0.001, ****p≤0.0001, and ns (not significant) p>0.05.

Also, Western blot analysis of CDK1 gene revealed that the protein fold change was affected by AURKA knockdown, as shown in Figure 4D, as it decreased to 0.24-fold compared to the control with a P-value ≤0.01.

The analysis of cell cycle phases by flow cytometry for HepG2 cells treated with both gRNA1 and gRNA2 revealed that the cells were significantly arrested in the G2 phase, as shown in Figure 4F, with 24.5 percent compared to the control with 14.7 percent (P-value ≤0.01).Additionally, HepG2-treated cells revealed a 54.69 percent decline in G1 compared to the control cells’ 60.96 percent (p-value ≤0.05), and a slight reduction in S phase compared to the control cells’ 20.13 and 22.23 percent, respectively. This pattern suggests an accumulation of cells at G2 phase rather than a general shift in the cell cycle.

These findings suggest that AURKA knockdown can regress HCC by enhancing the G2 cell cycle arrest, which confirms the previously reported involvement of AURKA in the G2 to M transition[60] and its enrichment in the G2M checkpoint hallmark gene set, as we showed in this study by the bioinformatics analysis (Figure 2).

### Knockdown of AURKA enhances apoptosis and impairs the proliferation of HepG2 cells

HepG2 apoptosis and its regulating genes, in addition to clonogenic potential, were examined to assess the effect of AURKA knockdown with gRNA1 and gRNA2 on HepG2 cell death and proliferation. The P38 and BCL-2 genes exhibited significant downregulation, with relative gene expression of 0.25 and 0.42-fold, respectively compared to the control, as shown in Figure 5A, B. However, Caspase8 and P53 genes showed significant upregulation with relative gene expression of 11.74-fold and 1.28-fold, respectively (p-value ≤0.01) compared to the control, as shown in Figure 5C, B.

**Figure 5:**
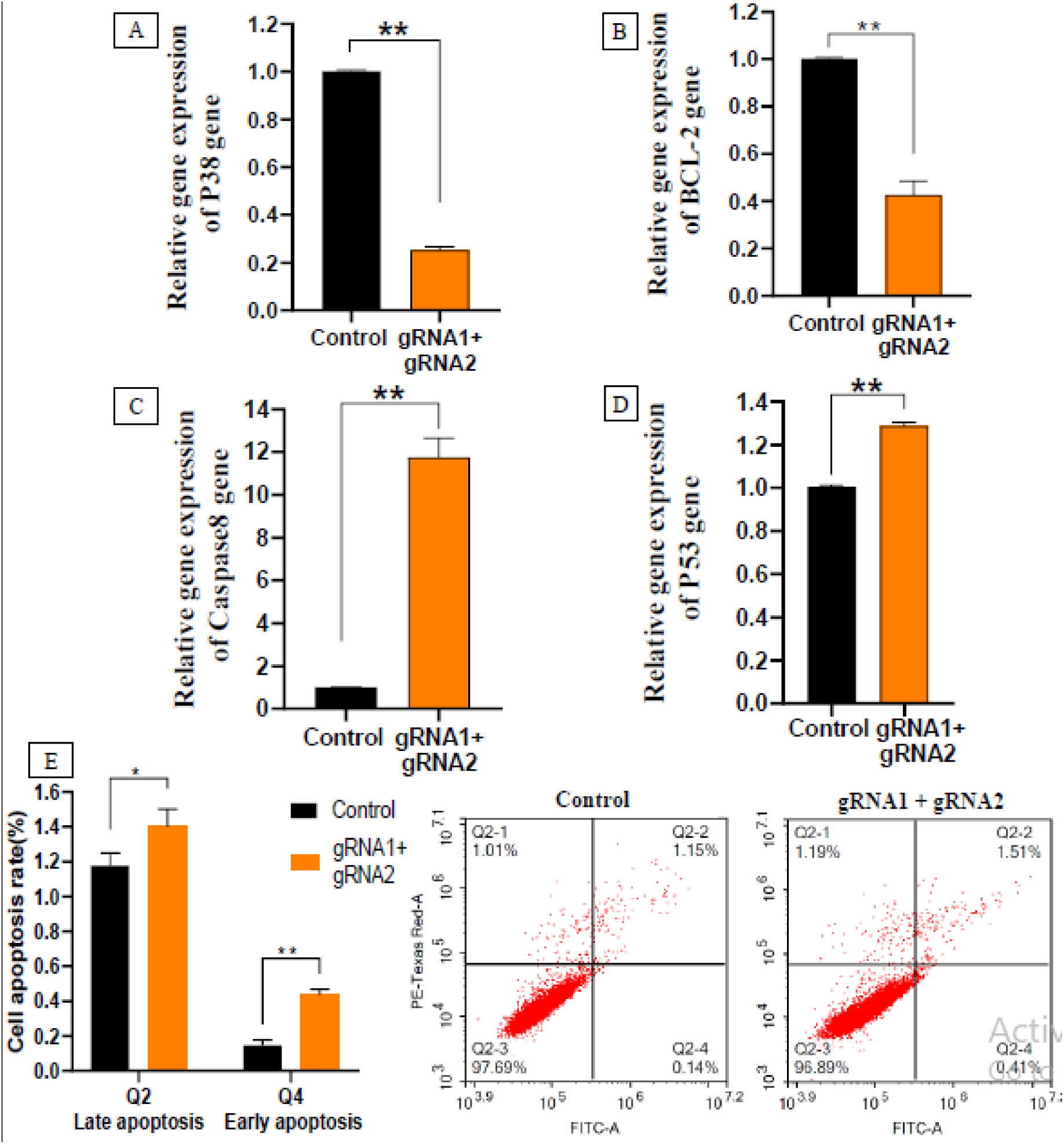
AURKA downregulation forced HepG2 cells going through apoptosis. AURKA downregulation impact on the relative expression of several apoptotic genes in Control (Null vector) and gRNA1+gRNA2 (HepG2 cells treated with both gRNA1 and gRNA2): A) P38, B) BCL-2, C) Caspase-8, D) P53. E) Apoptotic phase analysis in Control and gRNA1+gRNA2. All results were normalized to GAPDH. All experiments were performed as three independent experiments. GraphPad Prism version 8.0.2 was used to code, tabulate, and statistically analyse the collected data using unpaired t-tests or two-way ANOVA, and the Tukey-Kramer multiple comparisons test was used in the statistical study to compare the differences between the groups. The data are presented as the mean ± standard deviation (SD). Significant differences are indicated by the symbols *p≤0.05, **p≤0.01, ***p≤0.001, ****p≤0.0001, and ns (not significant) p>0.05.

The analysis of apoptosis by flow cytometry revealed that treated HepG2 cells significantly entered early apoptosis with 0.436 percentage compared to control cells of 0.146 cell percentage (p-value ≤0.01). The percentage of late apoptosis is also significantly higher in treated HepG2 cells equal 1.4 compared to the control 1.17 (P-value ≤0.05).

HepG2 cell growth was greatly impacted with AURKA knockdown. HepG2 cells showed a significant reduction in cell proliferation after transfection with gRNA1 and gRNA2 compared to the control with a p-value of ≤0.0001, as shown in Figure 6B. As demonstrated by these results, AURKA knockdown significantly regulated the pro- and anti-apoptotic genes, causing HepG2 cells to enter programmed cell death. Moreover, the anti-apoptotic effect of AURKA knockdown is complemented by reducing the proliferation of the HCC cells, where G2 accumulation is suggest here to be a primary mechanism by which gRNA1+gRNA2 treatment reduces proliferation, with apoptosis playing a minimal role.

**Figure 6:**
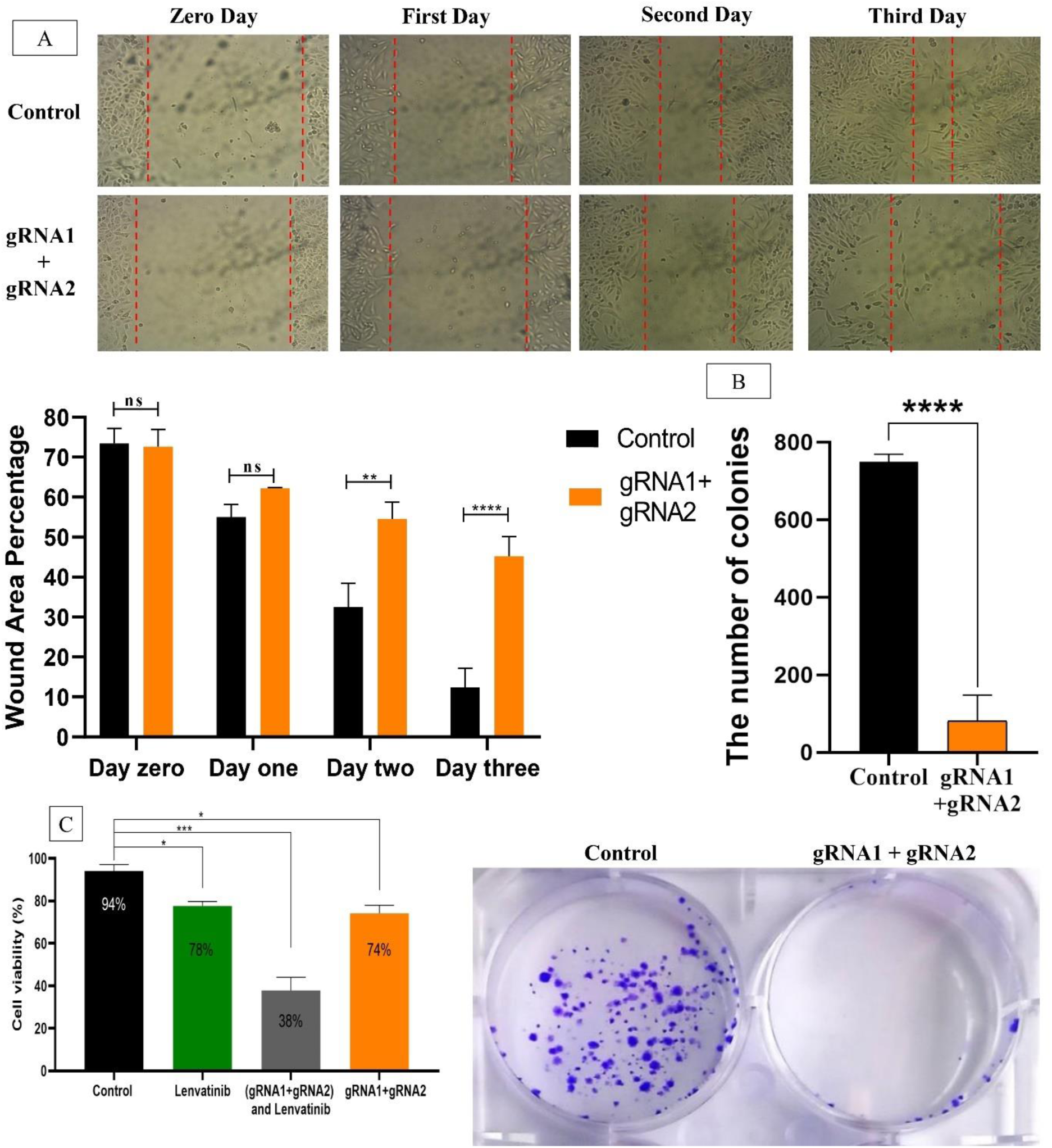
Knockdown of AURKA suppresses cell migration, colony formation, and viability in HCC cells. (A) Wound healing assay showing the effect of dual gRNA-mediated gene knockdown in HepG2 cells treated with both gRNA1 and gRNA2 (gRNA1 + gRNA2) compared to Control (Null vector) on cell migration at 0, 1, 2, and 3 days. Wound boundaries are indicated by red dashed lines in the representative photos. (B) Colony formation assay comparing the number of colonies formed in Control and gRNA1+gRNA2. (C) Cell viability assay showing the effect of Lenvatinib, gRNA1+gRNA2, and their combination on HCC cell survival. All results were normalized to GAPDH. All experiments were performed as three independent experiments. GraphPad Prism version 8.0.2 was used to code, tabulate, and statistically analyse the collected data using unpaired t-tests or ordinary one-way ANOVA, or two-way ANOVA, and the Tukey-Kramer multiple comparisons test was used in the statistical study to compare the differences between the groups. The data are presented as the mean ± standard deviation (SD). Significant differences are indicated by the symbols *p≤0.05, **p≤0.01, ***p≤0.001, ****p≤0.0001, and ns (not significant) p>0.05.

### Knockdown of AURKA delays HepG2 migration

AURKA gene knockdown has a major impact on numerous genes involved in cell migration, such as CDK1 and CDK2 [61]. HepG2 cell migration was substantially halted by the treatment with both gRNA1 and gRNA2 starting from 24 hours post-scratching, as seen in Figure 6A. After 48 h and 72 h of performing the scratch, the treated HepG2 cells showed a significant inhibition of cell migration with 54.4 % of wound area (P-value ≤0.01) and 45.2 % of wound area (P-value ≤0.0001), respectively.

These results indicated that AURKA knockdown not only impairs HCC cell proliferation and viability but is also able to reverse liver cancer cell migration characteristics.

### Knockdown of AURKA and Lenvatinib enhanced the inhibition of HepG2 cell viability

Lenvatinib exhibits notable benefits in HCC [62]. In our study, Lenvatinib caused 78% of HepG2 cell viability with a p-value ≤0.05 compared with the control. Similarly, the downregulation of AURKA caused a 74% of HepG2 cell viability with a p-value ≤0.05. As shown in Figure 6C, the combination of Lenvatinib and both sgRNA1 and sgRNA2 caused a more significant reduction in HepG2 cell viability by 37% with a P-value ≤0.001. Although, Bliss score here (0.1972, single dose, single time point) is exploratory and does not constitute evidence of pharmacological synergy, the viability analysis demonstrated that the combination of Lenvatinib and both gRNA1 and gRNA2 (gRNA1+gRNA2) had an enhanced effect.. These results indicated that combinations of targeted therapy with gene editing significantly improved the responsiveness of HCC..

## Discussion

One of the leading causes of cancer-related deaths globally is HCC, a widespread malignant tumor [63]. HCC is a genetically heterogeneous malignancy in which single-gene mutation play a significant role in the tumor’s growth, metastasis, and resistance to treatment [64]. Results for advanced HCC are still disappointing despite recent therapeutic improvements, which underscore the need for innovative treatments[24]. The potential of CRISPR-based approaches to provide a range of anti-cancer benefits has been revealed by preclinical studies employing HCC models [65]. *In vitro* research has shown promising results, such as apoptosis induction, invasion reduction, decreased migration, and inhibition of HCC cell proliferation [66]. By using SaCas9 in this study, we could reduce the expression of AURKA, as one of the prominent oncogenes in HCC, and consequently enhance apoptosis and induce cell cycle arrest.

AURKA is an oncogenic tumor biomarker that is upregulated in different types of cancer [67]. As our bioinformatics analysis demonstrated here, AURKA is overexpressed in HCC, Chen, C., et al.[15] and Zhang et al [68] proved that AURKA overexpression induces the epithelial–mesenchymal transition (EMT) and facilitates phenotype progression of HCC via the PI3K and AKT signalling pathways, thus stimulating tumor metastasis. AURKA expression is linked to cell-cycle regulation since it co-expresses with essential mitotic proteins and oncogenes, highlighting its function in tumour growth[60]. Furthermore, immune profiling reveals that elevated AURKA levels associate with tumour microenvironment, characterisedwith positive associations with immune-checkpoint molecules and negative correlations with cytotoxic T-cell profiles a pattern that could indicate immunological suppression.

Du R et al [69] reported that AURKA plays a role in immune evasion and may complicate responses to immune checkpoint medications, as indicated by trends of lower responsiveness to such treatments in high-AURKA groups. Overall, overexpression of AURKA appears to increase tumour aggressiveness and promote immune escape mechanisms. Moreover, inhibition of AURKA expression decreased the cell proliferation of liver cancer cells and induced apoptosis [67]. Also, cancer and other disorders were known to arise and progress as a result of deregulation of this AURKA non-catalytic activity[70]. Therefore, by using the CRISPR/SaCas9 vector, we could decrease the expression and functionality of the AURKA gene in HepG2 cells. This study’s results demonstrate how the use of two targeting sgRNAs altered the coding sequence of the AURKA gene by causing several random indels and resulted in a decrease in the relative expression of the AURKA gene in the treated HepG2 cells.

Aurora kinase A consists of an N-terminal domain (1–133aa), a kinase domain (134–383 aa), and a C-terminal domain (384-402 aa) [71]. The active conformation of the Aurora kinase domain is attributed to a hinge region connecting the β-stranded N-terminal lobe and the α-helical C-terminal lobe. Following the folding of the N- and C-terminal domains, the active site of Aurora kinases takes on a certain shape. Interestingly, certain hydrogen interactions connect the hinge of the AURKA kinase domain to the purine ring of adenosine[72]. As a result of the mutations formed here, the protein-coding sequence changed to give different amino acids which may lead to a disturbance in the configuration of the AURKA protein. Both gRNAs caused different nucleotide substitutions in the N-terminal domain of AURKA, which is mainly involved in protein-protein interactions[59]. Those nucleotide substitutions changed the amino acids as follows: V65V, K69K, P70L. According to UniProt (O14965 · AURKA_HUMAN), KLVSSHK[Ubiquitinylation]PVQNQK is a unique peptide as it contains two lysine amino acids (K). A protein’s stability, position within the cell, and ability to interact with other proteins can all be altered by SUMOylation, which occurs at the sites K63 and K69[73]. Moreover, gRNA2 caused a substitution of amino acid V 39 in a unique peptide (VTQQFPCQNPLPVNSGQAQRVL) (Variant ID: rs1315333827). Hence, these nucleotide changes may significantly influence the molecular mechanisms of AURKA transcription [74].

AURKA has been classified as a ‘G2/M’ gene [75]. The expression of AURKA varies depending on the stage of the cell cycle[74]. Several cell cycle checkpoints, such as the G1, spindle assembly, and G2, are disrupted by AURKA overexpression [76]. In this study, as a result of the knockdown of the AURKA gene by both gRNAs, the relative gene expression of p21 was significantly upregulated. As a cell cycle checkpoint, p21 can halt the progression of the cell cycle at the G1/S and G2/M transitions by inhibiting CDKs activity during specific phases of the cell cycle, such as CDK2 during the G1/S phase and CDK1 during the G2/M phase, resulting in growth arrest during those particular cell cycle phases[77]. Moreover, PI3K is phosphorylated by AURKA leading to increased expression of genes related to proliferation in HCC [78]. Our findings were validated by showing significant downregulation of CDK1, CDK2, and PI3K [79], followed by a consequent G2/M cell cycle arrest [80]. We also hypothesis that cell cycle arrest at the G2/M phase transition here as a result of blocking AURKA function might be independent of its kinase catalytic activity [81].

There is substantial evidence supporting the processes of AURKA-mediated carcinogenesis [71]. AURKA promotes cell proliferation, survival, and resistance to therapy in tumor cells by interacting with alternative oncogenic pathways such as MYC[82], p38/MAPK[83], and p53 [84] signalling pathways. Additionally, to provide tumor cells a survival advantage, AURKA alters pro-apoptotic (BCL-2, MLC1) expression levels [85]. The p38/MAPK signalling is crucial to cell proliferation and programmed cell death regulation for promoting cancer progression [86]. Tan, J., et al.[83] proved that the inhibition of AURKA is accompanied by inhibition of p38 /MAPK phosphorylation and inactivation of p38 /MAPK signalling leading to reduced expression of the anti-apoptotic protein (Bcl-2). Our bioinformatics and expression analyses here go strongly in line with the previous studies.

Aurora kinases control p53 activity by post-translationally altering the p53 protein directly or indirectly [84]. When AURKA is overexpressed, p53 is more readily degraded, which downregulates checkpoint-response pathways and promotes cell carcinogenesis[87]. The inverse relationship between p53 and AURKA has significant implications for anticancer treatment [59]. The previous correlations go in line with our data indicating that knocking down the AURKA gene increases the relative expression of the p53 gene. Caspase-8, a key player in numerous signalling pathways, is essential for controlling cytokine production, immune cell homeostasis, and programmed cell death [88]. A common finding in HCC is the deregulation of caspase-8 expression, which results in functional abnormalities in both the tumor microenvironment (TME) and HCC cells, and this imbalance may have substantial consequences on the aggressiveness, progression, and drug resistance of HCC [89]. Since AURKA normally suppresses apoptosis to promote cell survival, there is a correlation between the downregulation of AURKA and the overexpression of caspase 8, and cell caspase-based pro-apoptotic pathways may be activated when this function is suppressed [90]. Our finding revealed that Caspases 8 was upregulated when the AURKA gene was knocked down. The previous correlations have been validated here after downregulation of AURKA that promoted the early apoptotic phase with a higher cell percentage than the late apoptosis of HCC cells. In addition, as a known mitotic regulator and carcinogenic driver, the anti-proliferative and long-term growth and survival inhibition effects of AURKA downregulation emphasised the vital role of AURKA in preserving the ability of HCC cells to proliferate [60, 91]. Targeting the cells with the highest clonogenic potential supports the idea that AURKA is a possible molecular target in hepatocellular carcinoma, with implications for lowering tumour recurrence and resistance [92].

AURKA overexpression renders cancer cells resistant to anti-cancer drugs, while AURKA knockdown increases the susceptibility of cancer cells to Taxol-induced apoptosis through a number of signalling pathways, according to Jian-Ming Sun et al [80] and Xiaoting Chen et al [90]. One new first-line treatment for hepatocellular carcinoma (HCC) is Lenvatinib, a multi-target tyrosine kinase inhibitor (TKI)[93]. Lenvatinib (multitargeted inhibitor) suppresses vascular endothelial growth factor receptor (VEGFR) 1–3, fibroblast growth factor receptor (FGFR) 1–4, platelet-derived growth factor receptor (PDGFR) α, as well as proto-oncogenes RET (rearranged during transfection) and KIT (stem cell factor receptor)[94]. Numerous preclinical cancer models demonstrate Lenvatinib’s anticancer and immunomodulatory efficacy particularly in HCC[95]. Our research has demonstrated that AURKA knockdown enhanced HepG2 cellsresponsiveness by reducing their viability after Lenvatinib treatment. Our findings showed that the cell viability of HepG2 cells treated with both Lenvatinib and gRNA1 and gRNA2 (gRNA1+gRNA2) was below 40% when compared to the untreated cells and the individual treatment, and this combination had a favorable combined effect. Also, the effect of the AURKA gene knocked down with only both gRNA1 and gRNA2 (gRNA1+gRNA2) was slightly less than when Lenvatinib was used alone. Moreover, Lenvatinib resistance is promoted by AURKA’s work, Abudoureyimu, M., et al[96] indicate that decreased sensitivity to Lenvatinib is associated with increased AURKA expression, suggesting that AURKA knockdown may cause cells to become more sensitive. Han, K., Jeng, E., Hess, G. et al. [97] supported the potential of combining targeted therapy and gene-based interventions for improved therapeutic outcomes.Together, these findings provide credence to the idea that AURKA has a role in HCC survival and therapy resistance and that combination targeting could result in improved HCC treatment success.

## Conclusion

Our *In silico* findings demonstrated that AURKA overexpression plays a pivotal role in HCC samples. Our knockdown results demonstrate that AURKA depletion suppressed HCC progression by inducing cell cycle arrest, decreasing cell proliferation, reducing apoptosis, and altering key signalling pathways. CRISPR/SaCas9-mediated knockdown of AURKA significantly reduced its gene and protein expression, leading to cell cycle arrest, increased pro-apoptotic genes, and reduced cancer development potential. This anti-tumor effect was shown through enhanced expression of tumor suppressor genes, while decreasing oncogenic genes, as well as deteriorating the functionality of HCC cells. Furthermore, combining AURKA knockdown with Lenvatinib treatment yielded an enhanced anti-cancer effect. These *in vitro* molecular and functional anti-HCC benefits of AURKA knockdown by gene-editing approach provide advantages of AURKA as a promising molecular target for prospective additional in vivo and translational research. Future *in vivo* experiments and clinical implementations are required to validate our suggested strategy to target HCC.

## Strengths and Limitations

The coupling of thorough experimental validation with bioinformatics-driven target discovery from human-derived datasets, selected based on clearly defined criteria to ensure both biological relevance and technical robustness for differential expression analysis, strengthens this study and supports AURKA as a pertinent therapeutic target in hepatocellular carcinoma (HCC). The CRISPR/SaCas9 technique guarantees accurate AURKA knockdown. The experimental design uses flow cytometry, functional tests, and molecular studies to evaluate different cellular characteristics and carcinogenic pathways. Furthermore, the combination treatment with Lenvatinib raises the possibility to enhance therapeutic, and clinical utility. Notwithstanding its advantages, the study contains some drawbacks. It mostly relies on a single HCC cell line (HepG2), which may overlook disease heterogeneity. The absence of *in vivo* validation limits the evaluation of therapeutic efficacy and safety. Additionally, while its interaction with Lenvatinib appears promising, further investigation is required across different models, dosing, timing, and delivery strategies.

## Recommendation and future perspectives

Further studies should be conducted to validate the therapeutic potential of AURKA targeting in hepatocellular carcinoma (HCC) utilizing more representative models, including multiple cell lines, patient-derived systems, and *in vivo* approaches to assess efficacy, safety, and pharmacokinetics. Optimizing CRISPR/SaCas9 *in vivo* delivery methods, including HCC-targeted viral and non-viral systems, and long post-therapeutic observation should be the main goals of future research to ensure the maximum ability to deliver the gene-editing cargo to all of the cells within a tumor and to avoid development of “escape” mutations that would presumably drive the re-emergence leading to tumor recurrence. Furthermore, to develop successful combination strategies, the enhancement seen with Lenvatinib needs to be further investigated through dose-response analysis and preclinical validation. The goal of these recommendations is to use AURKA-targeted CRISPR techniques in effective HCC gene therapy.

## Acknowledgment

We want to express our gratitude to Professor Dr. Ahmed Ragab Hamed (Chemistry of Medicinal Plants Department and Biology Unit, Central Laboratory for Pharmaceutical and Drug Industries Research Institute, National Research Centre, Cairo, Egypt) for his valuable comments on the Western blot technique. We are so grateful to Dr.Shaymaa M.M.Yahya (Hormones Department, Medicine and Clinical Studies Research Institute, and Stem Cell Lab, Centre of Excellence for Advanced Sciences, National Research Centre, Cairo, Egypt) for her contribution to the colony formation assay and for providing us with the primers of CDK1, Caspase 8, BCL2, P53, and P21 genes to perform the RT-PCR assay. We are thankful to Associate Professor Dr. Asmaa M. Elfiky (Environmental and Occupational Medicine Department, National Research Centre, Cairo, Egypt) for providing us with the primers of CDK2, P38, and PI3K (alpha subunit) genes to perform the RT-PCR assay and for her fruitful comments on the MTT assay.

## Declarations

### Ethics approval and consent to participate

Not applicable.

### Clinical trial number

Not applicable.

### Consent for publication

Not applicable.

### Availability of data and material

The data that support the findings of this study are available upon reasonable request to the corresponding author.

### Authors’ contributions

N.S: Conceived the work, conducted the experiments and data analysis, wrote the manuscript, and created the figures.

M.E: Proposed the main idea, conceived the work, revised the manuscript, and supervised the work.

R.M: Put the study design and the methodology, supervised the work, and revised the manuscript.

E.A.M: Execution of the primary bioinformatics analysis execution of the primary that identified and selected AURKA as the therapeutic target for the HCC model

B.S: Supervision.

All authors have read and approved the final manuscript.

## Abbreviations

HCC: Hepatocellular carcinoma (HCC)
AURKA: Aurora kinase A
LIHC: Liver Hepatocellular Carcinoma
TCGA: The Cancer Genome Atlas
EMT: epithelial-mesenchymal transition
CSC: cancer stem cell
CRISPR: clustered regularly interspaced short palindrome repeats
SgRNA: single-guide RNA
SpCas9: Streptococcus pyogenes Cas9
SaCas9: Staphylococcus aureus Cas9
GEPIA: Gene Expression Profiling Interactive Analysis
DEGs: Differentially expressed genes
WGCNA: Weighted Gene Co-expression Network Analysis
ORA: over-representation analysis
TME: tumor microenvironment
TIDE: Tumor Immune Dysfunction and Exclusion
ICB: Immune Checkpoint Blockade
CIAP: calf intestinal alkaline phosphatase

---

Supplementary Figure 1: Principal Component Analysis (PCA) for the normal GTEx liver gene dataset and TCGA LIHC dataset

Supplementary Figure 2: Weighted Gene Co-expression Network Analysis (WGCNA)

Supplementary Figure 3: Expression of AURKA in TCGA-LIHC

Supplementary Table: TIDE Score

