## Supplementary figure 1 for "Targeting AURKA by CRISPR/SaCas9 Alters Cell Cycle Progression and Enhances Lenvatinib-Associated Reduction in Cell Viability in Hepatocellular Carcinoma: An Integrated In Silico and HepG2 in Vitro Study"

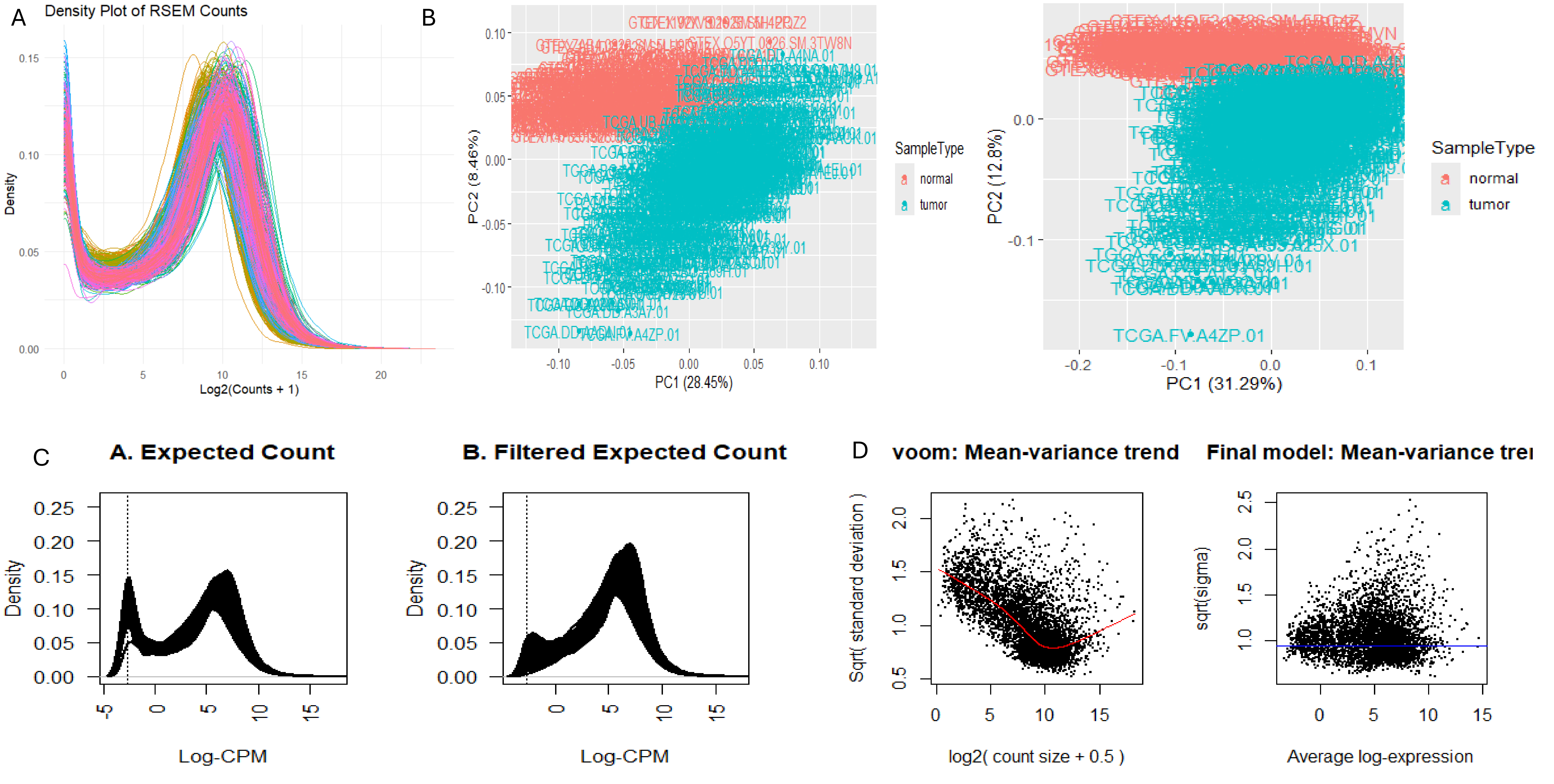

A
Density Plot of RSEM Counts
Density
Log2(Counts + 1)
B
PC2 (8.46%)
PC1 (28.45%)
SampleType
normal
tumor
PC2 (12.8%)
PC1 (31.29%)
TCGA.FV.A4ZP.01
C
A. Expected Count
B. Filtered Expected Count
Density
Log-CPM
D
voom: Mean-variance trend
Final model: Mean-variance tre
Sqrt( standard deviation )
log2( count size + 0.5 )
sqrt(sigma)
Average log-expression
