## supplementary figure 2 for "Targeting AURKA by CRISPR/SaCas9 Alters Cell Cycle Progression and Enhances Lenvatinib-Associated Reduction in Cell Viability in Hepatocellular Carcinoma: An Integrated In Silico and HepG2 in Vitro Study"

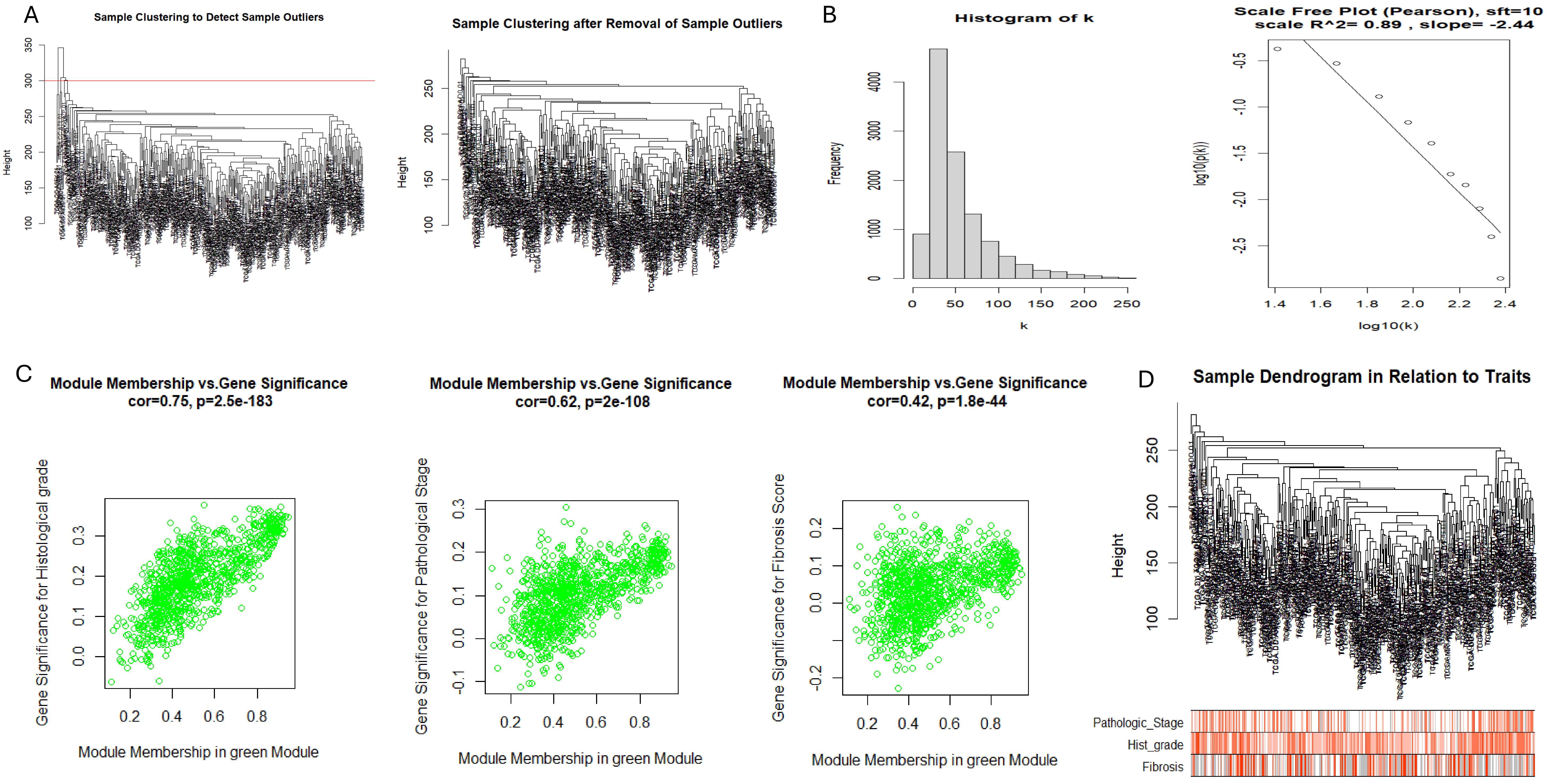

A
Sample Clustering to Detect Sample Outliers
Height
Sample Clustering after Removal of Sample Outliers
Height
B
Histogram of k
Frequency
k
Scale Free Plot (Pearson), sft=10
scale R^2= 0.89 , slope= -2.44
log10(p(k))
log10(k)
C
Module Membership vs.Gene Significance
cor=0.75, p=2.5e-183
Gene Significance for Histological grade
Module Membership in green Module
Module Membership vs.Gene Significance
cor=0.62, p=2e-108
Gene Significance for Pathological Stage
Module Membership in green Module
Module Membership vs.Gene Significance
cor=0.42, p=1.8e-44
Gene Significance for Fibrosis Score
Module Membership in green Module
D
Sample Dendrogram in Relation to Traits
Height
Pathologic_Stage
Hist_grade
Fibrosis
