## supplementary figure 3 for "Targeting AURKA by CRISPR/SaCas9 Alters Cell Cycle Progression and Enhances Lenvatinib-Associated Reduction in Cell Viability in Hepatocellular Carcinoma: An Integrated In Silico and HepG2 in Vitro Study"

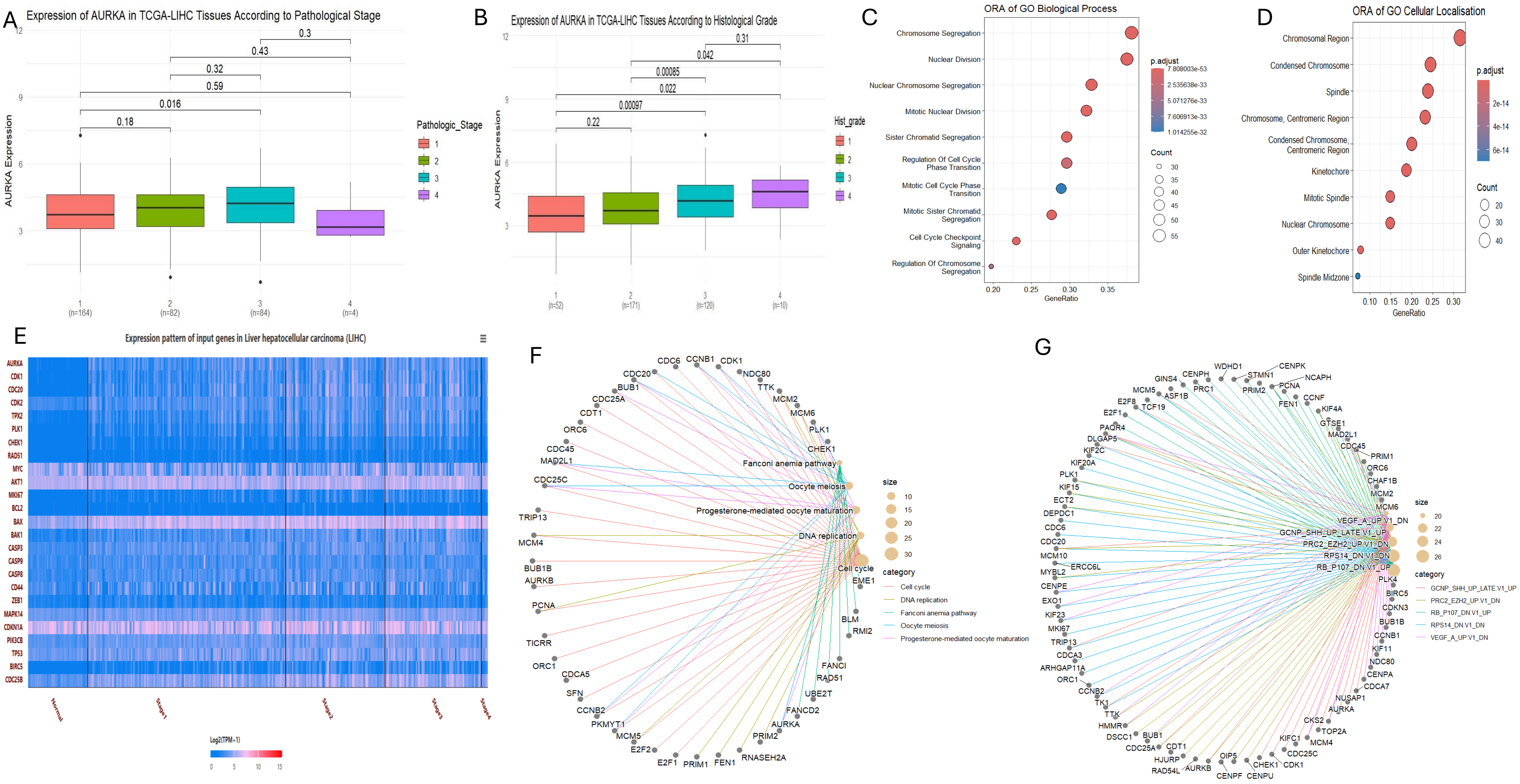

A
Expression of AURKA in TCGA-LIHC Tissues According to Pathological Stage
AURKA Expression
0.3
0.43
0.32
0.59
0.016
0.18
Pathologic_Stage
1
2
3
4
1 (n=164)
2 (n=82)
3 (n=84)
4 (n=4)
B
Expression of AURKA in TCGA-LIHC Tissues According to Histological Grade
AURKA Expression
0.31
0.042
0.00085
0.022
0.00097
0.22
Hist_grade
1 (n=52)
2 (n=171)
3 (n=120)
4 (n=10)
C
ORA of GO Biological Process
Chromosome Segregation
Nuclear Division
Nuclear Chromosome Segregation
Mitotic Nuclear Division
Sister Chromatid Segregation
Regulation Of Cell Cycle Phase Transition
Mitotic Cell Cycle Phase Transition
Mitotic Sister Chromatid Segregation
Cell Cycle Checkpoint Signaling
Regulation Of Chromosome Segregation
GeneRatio
p.adjust
7.808003e-53
2.535638e-33
5.071276e-33
7.606913e-33
1.014255e-32
Count
30
35
40
45
50
55
D
ORA of GO Cellular Localisation
Chromosomal Region
Condensed Chromosome
Spindle
Chromosome, Centromeric Region
Condensed Chromosome, Centromeric Region
Kinetochore
Mitotic Spindle
Nuclear Chromosome
Outer Kinetochore
Spindle Midzone
GeneRatio
p.adjust
2e-14
4e-14
6e-14
Count
20
30
40
E
Expression pattern of input genes in Liver hepatocellular carcinoma (LIHC)
AURKA
CDK1
CDC20
CDK2
TPX2
PLK1
CHEK1
RAD51
MYC
AKT1
MKI67
BCL2
BAX
BAK1
CASP3
CASP9
CASP8
CD44
ZEB1
MAPK14
CDKN1A
PIK3CB
TP53
BIRC5
CDC25B
Normal
Stage1
Stage2
Stage3
Stage4
Log2(TPM+1)
0
5
10
15
F
Fanconi anemia pathway
Oocyte meiosis
Progesterone-mediated oocyte maturation
DNA replication
Cell cycle
size
10
15
20
25
30
category
Cell cycle
DNA replication
Fanconi anemia pathway
Oocyte meiosis
Progesterone-mediated oocyte maturation
G
VEGF_A_UP.V1_DN
GCNP_SHH_UP_LATE.V1_UP
PRC2_EZH2_UP.V1_DN
RPS14_DN.V1_DN
RB_P107_DN.V1_UP
size
20
22
24
26
category
GCNP_SHH_UP_LATE.V1_UP
PRC2_EZH2_UP.V1_DN
RB_P107_DN.V1_UP
RPS14_DN.V1_DN
VEGF_A_UP.V1_DN
